# Silencing of Exosomal miR-181a reverses Pediatric Acute Lymphocytic Leukemia Cell Proliferation

**DOI:** 10.1101/2020.05.10.086967

**Authors:** Shabirul Haque, Sarah R. Vaiselbuh

## Abstract

Exosomes are cell-generated nano-vesicles (30-150 nm) found in most biological fluids. Major components of their cargo are lipids, proteins, RNA, DNA, and non-coding RNAs. Exosomes carry the fingerprint of the parental tumor and as such, may regulate tumor growth, progression and metastasis. We investigated the impact of exosomes on cell proliferation in pediatric acute lymphocytic leukemia and its reversal by silencing of exo-miR-181a.We isolated exosomes from serum of acute lymphocytic leukemia pediatric patients (Exo-PALL) and conditioned medium of leukemic cell lines (Exo-CM) by ultracentrifugation. Gene expression was carried out by q-PCR. We found that Exo-PALL promote cell proliferation in leukemic B cell lines as well as in the control B cell line. This exosome-induced cell proliferation is a precise event with up-regulation of proliferative (PCNA, Ki-67) and pro-survival genes (MCL-1, and BCL2), and suppression of pro-apoptotic genes (BAD, BAX). Exo-PALL and Exo-CM both show over expression of miR-181a compared to controls (Exo-HD). Specific silencing of *exosomal* miR-181a using a miR-181a inhibitor confirms that miR-181a inhibitor treatment reverses Exo-PALL/Exo-CM-induced leukemic cell proliferation *in vitro*. Altogether, this study suggests that exosomal miR-181a inhibition can be a novel target for growth suppression in pediatric lymphatic leukemia.

## Introduction

Exosomes are spherical nanoparticles (30-150 nm) produced by cells in most body fluids (i.e. serum, urine, breast milk, ascites) under normal and pathological conditions. Exosomes originate from the cytoplasm of the cell by inward budding of multivesicular bodies^1^.Consequently, exosomal content is enriched for proteins, lipids, cytoplasmic mRNA and most importantlynon-codingmicro-RNA^2-3^.Several investigators have established that biologically functional microRNAs (miRNAs) are transported by exosomes and exosomes-encapsulated miRNAs can be delivered to target cells. The exosomal cargo that is transferred as such is functionally active; enabling the target cells to reprogram and redefine their immunological responses at a molecular level ^4-6^.Moreover, exosomes derived from the tumor microenvironment can recruit and impair normal cells to contribute to the disease status. Some reports in literature support the fact that miRNAs carried within exosomes reveal real-time information regarding disease status in leukemia and other cancers, and therefore exosomes have been studied for their potential as novel biomarkers for cancer, easily accessible in the serum of patients by minimally invasive techniques such as a simple phlebotomy^7-10^. Our work focuses on the functional role of exosomal-miRNA-181a (exo-miR-181a) in pediatric acute lymphocytic leukemia (PALL).

MicroRNAs are endogenously expressed non-coding regulatory RNAs of 20-25 long nucleotide and play a major role as gene expression regulators. They can act as gene expression silencers at a post-transcriptional level either by inhibiting mRNA translation or by mRNA degradation. Furthermore, miRNAs can destabilize the mRNA transcript through binding to the 5’ or 3’ untranslated regions (UTR)^11-13^.As such, miRNA scan function as facilitators of fundamental normal biological processes such as cell differentiation, proliferation, apoptosis and survival by manipulation of target genes^14^.Dysregulation of these gene functions may be one of the contributing factors that can lead to leukemogenesis^15^. Precisely, the miR-181 family consists of four mature individual miRs: miR-181a, miR-181b, miR-181c, miR-181d that are highly conserved in evolution within almost all vertebrates, indicating their functional relevance^16^. Several investigators have established that miR-181a specifically modulates cellular events at multiple levels - such as cell proliferation, growth, survival and even chemo-sensitivity in cancer^17-18^. Yang et al. described that elevated expression of miR-181a leads to cancer progression and is associated with poor survival, suggesting clinical significance for the role of miR-181a^19-22^. In contrast, others have shown that down-regulated expression of miR-181a leads to poor prognosis, metastasis and cancer development^23-24^. This dual behavior of miR-181a has been reported in different types of cancer suggesting that the mechanism by which miR-181a exerts its functional role might be very disease-specific. There are only a few reports in literature that studied the role of *cellular* miR-181a in the proliferation of PALL. To the best of our knowledge, no one investigated so far the role of *exosomal* miR-181a in PALL progression.

Acute lymphoblastic leukemia (ALL) is one of the most common hematological cancers impacting both adults and children^25^. The peak age of pediatric ALL (PALL) incidence is between 2-4 years^26^. Some studies show that epigenetic factors, such as DNA methylation, gene regulation and chromatin remodeling^27^, play a significant role in PALL development and progression by dysregulation of miRNA^28-29^. Therefore, identification of unique miRNA expression patterns in PALL may have great potential for clinical translational therapy. We explored the role of PALL exosomes (Exo-PALL, derived from serum of children diagnosed with ALL or from conditioned medium (CM) of ALL cell lines) and its possible correlation to leukemic cell proliferation. Exo-PALL showed up-regulated miR-181a expression compared to exosomes-derived from healthy donors serum. Moreover, exosome-induced cell proliferation was reversible by silencing of Exo-miR-181a by a specific miR-181a inhibitor. Silencing of miRNA/mRNA at cellular level is widely accepted and most commonly performed in scientific communities, but silencing at lipid nanoparticle/exosomes level are not common yet, its emerging. A recent article demonstrated cellular gene (indoleamine2, 3-dioxygenase 1) silencing by lipid nanoparticle loaded with gene specific siRNA showed significant effect by downregulatingindoleamine2, 3-dioxygenase 1, which showed lipid nanoparticle is vehicle/delivery system for siRNA^46^.

## Results

### Confirmation of CD63 and CD81 expression on exosomes by flow cytometer

Without any dispute, well established exosome biomarkers are CD63 and CD81 expression on the surface of exosomes. We explored expression of CD63 and CD81 on exosomes derived from serum and conditioned medium of cell line. Exosomes were captured with CD63 antibody coated magnetic beads overnight at room temperature. Then CD63 antibody captured exosomes were stained with CD81-PE conjugated antibody in including possible controls. Each treated conditions were analyzed by flow cytometer. Analyzed data demonstrate that exosomes are positive for CD81 and CD63 expression **(Fig. 1)**. We have shown in our previous published article that exosomes are positive with CD63 and negative with calnexin by western blot. We also examined and confirmed the size of the exosomes by nano tracking analysis (NTA-Malvern) which showed 90% of the exosomes are within the optimum size range of a 50-100 nm diameter^32^.

**Figure 1.**
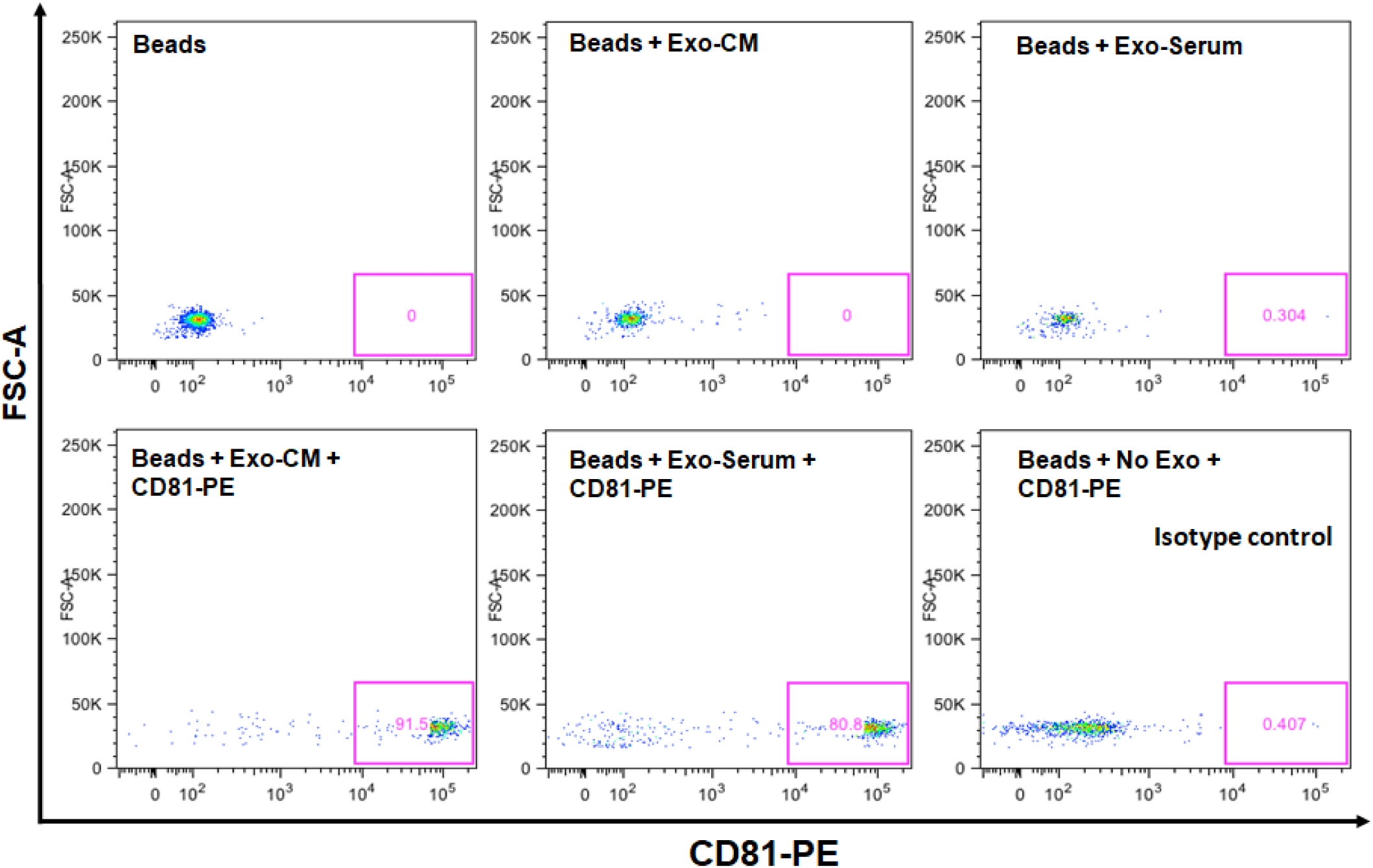
Expression of CD63 and CD81 on exosomes by flow cytometer. Binding/capture of exosomes was carried out with CD63 antibody coated magnetic beads for overnight at room temperature. Beads captured exosomes were stained with primary antibody CD81 antibody (Biotin conjugated), then stained with secondary detection reagent (streptavidin-PE conjugated). Each condition is labeled for control purpose in the figure. Exo-serum represents, exosomes derived from serum. Exo-CM represents, exosomes derived from conditioned medium of JM1 cell.

### Exo-JM1/SUP-B15 and Exo-PALL induces cell proliferation in cell lines

Exosomes were isolated from conditioned media (Exo-CM) of two ALL cell lines (JM1 and SUP-B15) and one control B cell line (CL-01). In addition, exosomes were harvested from human serum samples of either patients with childhood acute lymphocytic leukemia (Exo-PALL) or healthy donors (Exo-HD). In order to establish exosome dose titration and time kinetics of the process, Exo-CM (CL-01 and SUP-B15) was added to JM1 cells (paracrine incubation) in one of three different concentrations (100, 250, 500 µg/ml). PBS only was added as negative control. Cell counting was carried out at 24 and 48 hours under light microscopy. Control Exo-CL-01 did not induce cell proliferation at either time points, while Exo-SUP-B15 significantly induced cell proliferation compared to PBS control both at 24 and 48 hours. Similar results were obtained with autologous incubation (autocrine fashion) of Exo-JM1 on JM1 cells (data not shown). Exosome treatment augmented cell proliferation at a starting exosome concentration (100 µg/ml), with optimal proliferation at 250 µg/ml **(*Supplementary Fig. 1A)***. Consequently, we chose an exosome concentration of 250 µg/ml (noted saturated cell proliferation induction) as the working dose for future experiments. Similarly, we optimized the dose and time points from HD and PALL-derived serum exosomes. JM1 cells were treated with Exo-HD and Exo-PALL. The control Exo-HD did not induce cell proliferation compared to PBS at either time points (24 or 48 hours), while Exo-PALL significantly induced cell proliferation compared to Exo-HD as well as PBS control both at 24 and 48 hours **(*Supplementary Fig. S1-B)***. Based on these, we decided that 250 µg/ml exosome concentration and 24 hours incubation are the optimum conditions for the cell proliferation assay and we used these in the subsequent experiments.

Next step, we confirmed that CM-exosomes induce cell proliferation both in a paracrine or autocrine fashion using Exo-JM1 **(Fig. 2A)** and Exo-SUP-B15 **(Fig. 2B)**. Collectively, our data showed that exosomes originated from ALL cell lines (JM1, SUP-B15) promoted cellular proliferation in control human B cells (CL-01) as well as both leukemic B cell lines **(Figs. 2A and 2B)**. In addition, we wanted to explore the reproducibility of this cell proliferation effect with exosomes isolated from human serum from PALL patients (n=6) **(Figs. 2C and 2D)**. Exo-PALL at day 1 of diagnosis (Patient # 5) and Exo-HD from a healthy donor (HD# 1) were incubated with control human B cells (CL-01) and leukemia cell lines (JM1 and SUP-B15). Only P-ALL serum-derived exosomes promoted cell proliferation in control human B cells (CL-01) as well as leukemic B cell lines (JM1, SUP-B15) **(Fig. 2C)**. To consolidate our data, JM1 cells were treated with Exo-PALL from five other patients (Pt# 1, 2, 3, 4, and 6). All five Exo-PALL enabled significant induction of cell proliferation in JM1 cells **(Fig. 2D)**. Apart from cell counting using trypan blue, we also confirmed our cell proliferation experiment by MTS assay ***(Supplementary Fig. 2)***, supporting that treatment with leukemia derived exosomes induces augmented cell proliferation.

**Figure 2.**
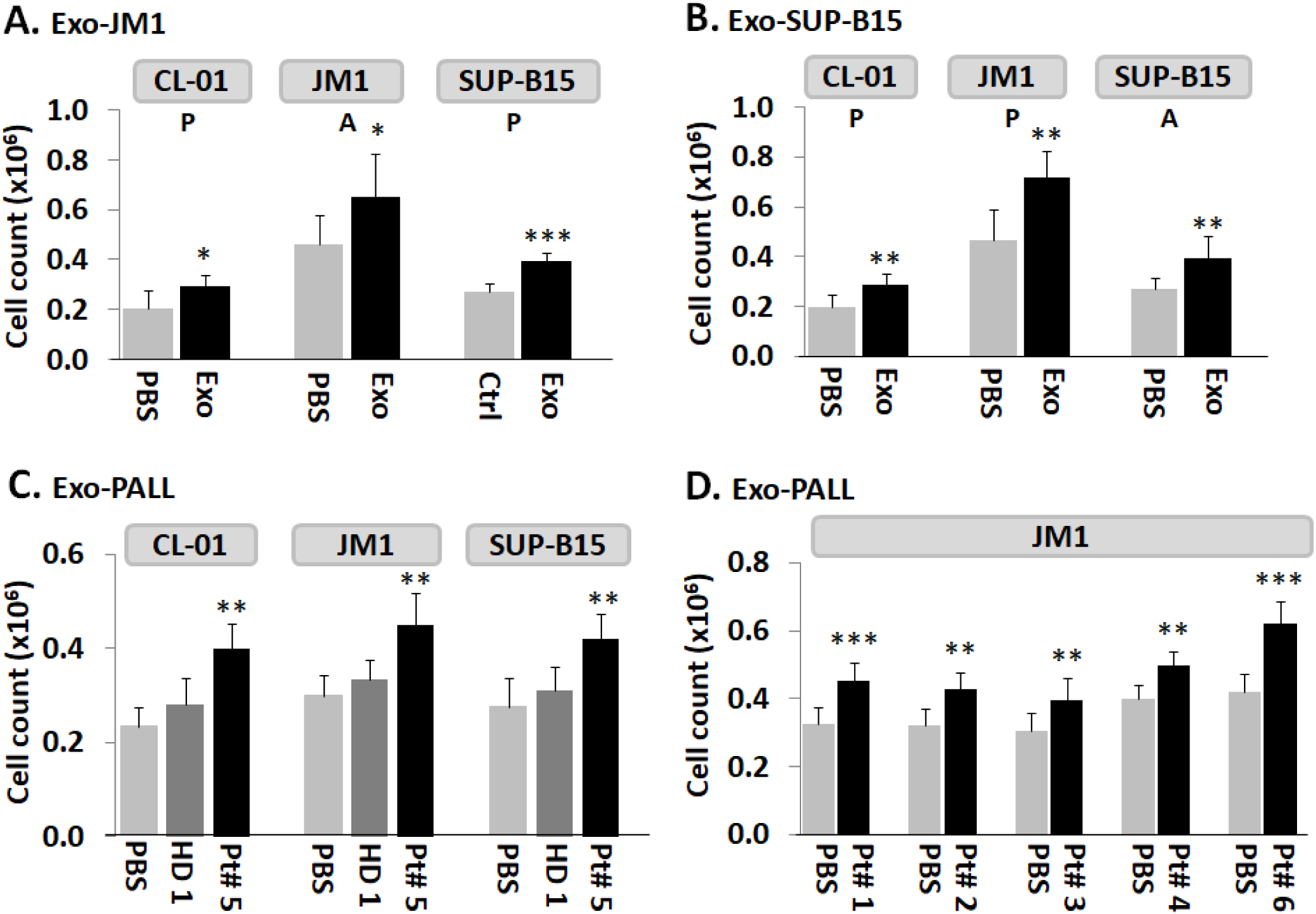
Exosomes from leukemia cell lines and from serum of pediatric ALL patients induce autocrine and paracrine cell proliferation in leukemia cell lines. **(A)** Exo-JM1-induced cell proliferation in control B cell line CL-01 and human leukemia cell lines JM1 and SUP-B15. **(B)** Exo-SUP-B15 - induced cell proliferation in control B cell line CL-01 and human leukemia cell lines JM1 and SUP-B15. **(C)** Exosomes derived from human serum of ALL patient#5 (Exo-PALL) at day 1 diagnosis induce proliferation in CL-01, JM1 and SUP-B15 cells compared to healthy donor exosomes (HD) or control PBS treated only. **(D)** Human serum exosomes (Exo-PALL) derived from five different patients (Pt # 1, 2, 3, 4 and 6) induce cell proliferation in JM1 cells compared to control PBS group.*(PBS: control - no exosomes; P: paracrine effect; A: autocrine effect. Mean of 3 experiments. P value *p<0.05, **p<0.01, ***p<0.001)*

### Exo-JM1 and Exo-SUP-B15 treatment augments proliferative/pro-survival genes, and down regulates pro-apoptotic genes

In order to evaluate one of the molecular pathways by which Exo-JM1 and Exo-SUP-B15 promotes cell survival and cell proliferation, we analyzed gene expression profiles in the JM1 leukemia cell line after exosomes exposure. Both Exo-JM1 and Exo-SUP-B15 altered the mRNA expression of regulatory genes such as Ki-67, BCL2, BAD in JM1 cells **(Figs. 3A-B)**.

**Figure 3.**
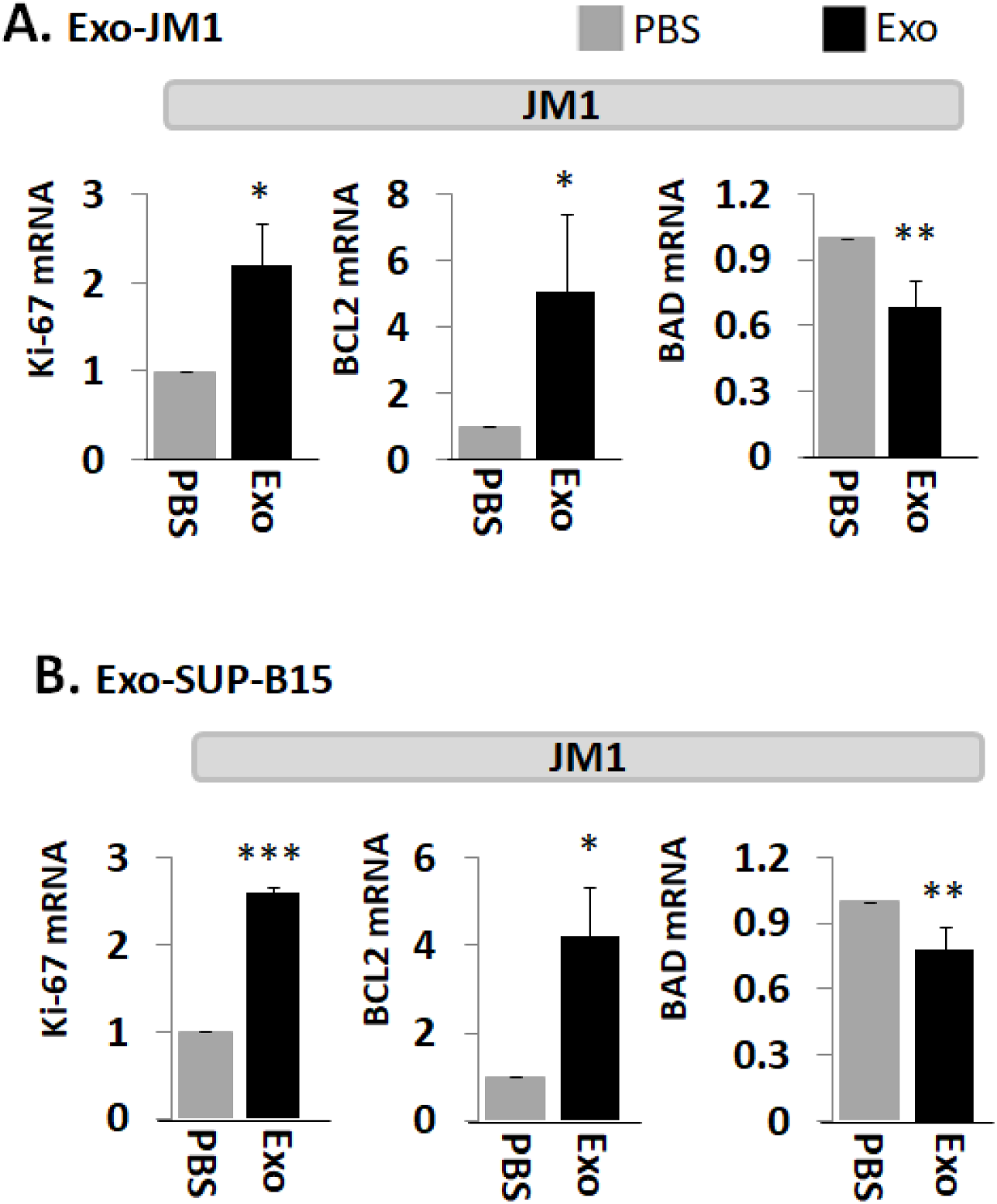
Exo-JM1 and Exo-SUP-B15 regulate proliferative, pro-survival, and pro-apoptotic genes in JM1 cells. **(A)** Exo-JM1 up-regulates Ki-67, and BCL2 mRNA expression (pro-survival) and down-regulates BAD mRNA expression (pro-apoptotic) in autocrine manner. **(B)** Exo-SUP-B15 promotes Ki-67 and BCL2 mRNA expression and down regulates BAD expression in JM1 cells in paracrine manner. Data represented are mean of three experiments. *(Ctrl: PBS only – no exosomes. P value * p<0.05, **p<0.01, ***p<0.001)*

Exo-CM from ALL cell lines (JM1 and SUP-B15) was added to JM1 cells and mRNA expression of proliferative (Ki-67, PCNA), pro-survival (MCL1, BCL2) and pro-apoptotic (BAD, BAX) genes was analyzed at two time points (6 and 24 hours). For both time points, incubation of JM-1 cells with Exo-JM1 (autocrine exposure) resulted in up-regulated expression of the proliferative/pro-survival genes Ki-67, PCNA, MCL1, and BCL2 while down-regulated expression of pro-apoptotic genes BAD and BAX ***(Supplementary Fig. 3A)***. To authenticate these changes in gene expression in cells that were exposed to exosomes, we also tested Exo-SUP-B15 on JM1 cells (paracrine exposure). We found a similar pattern of augmented expression of Ki-67, PCNA, MCL1, and BCL2 mRNA and down-regulated expression of BAD and BAX mRNA at both 6 and 24 hours induced by Exo-SUP-B15 ***(Supplementary Fig. 3B)***. Moving forward, we selected an incubation time of 24 hrs to replicate the results in other experiments. Overall, our current data suggest that both Exo-JM1 and Exo-SUP-B15 promote cell proliferation by enhancing proliferative/pro-survival signals and suppressing pro-apoptotic genes.

### Exo-PALL promotes proliferative/pro-survival genes, and down-regulates pro-apoptotic genes

Next, we tested the effect of exosomes of serum samples from ALL patients and healthy donors on gene induction **(Fig. 4)**. Exosomes were co-cultured with three leukemia cell lines (JM1, SUP-B15, and NALM-6). To confirm the results obtained with Exo-CM, we explored if Exo-PALL induced the same changes in gene expression involved in the proliferation, survival and apoptotic cellular pathways. We used exosomes from serum of patients at different stages of leukemia (at diagnosis, first remission, relapse and second remission). Exo-PALL at the diagnosis (Day 1) augmented Ki-67 and BCL2 while BAD gene expression appeared to be down-regulated **(Fig. 4A)**. Interestingly, Exo-PALL from Day 29 (1^st^ remission) failed to induce Ki-67 and BCL2 in target cells, while the BAD gene was up regulated **(Fig. 4A)**. This experiment was carried out in three different target leukemia cell lines. We also compared the potential of human serum exosomes isolated from PALL relapse versus PALL 2^nd^ remission (pt #3 and pt #4). PALL relapse exosomes exposure up-regulated Ki-67, BCL2 and down regulated BAD mRNA expression **(Fig. 4B)**. In contrast, incubation of target cells with PALL 2^nd^ remission-derived exosomes failed to up-regulate Ki-67 and BCL2 mRNA and failed to down-regulate BAD mRNA expression **(Fig. 4B)**. These experiments were also carried out in three different target leukemia cell lines with consistent results obtained.

**Figure 4.**
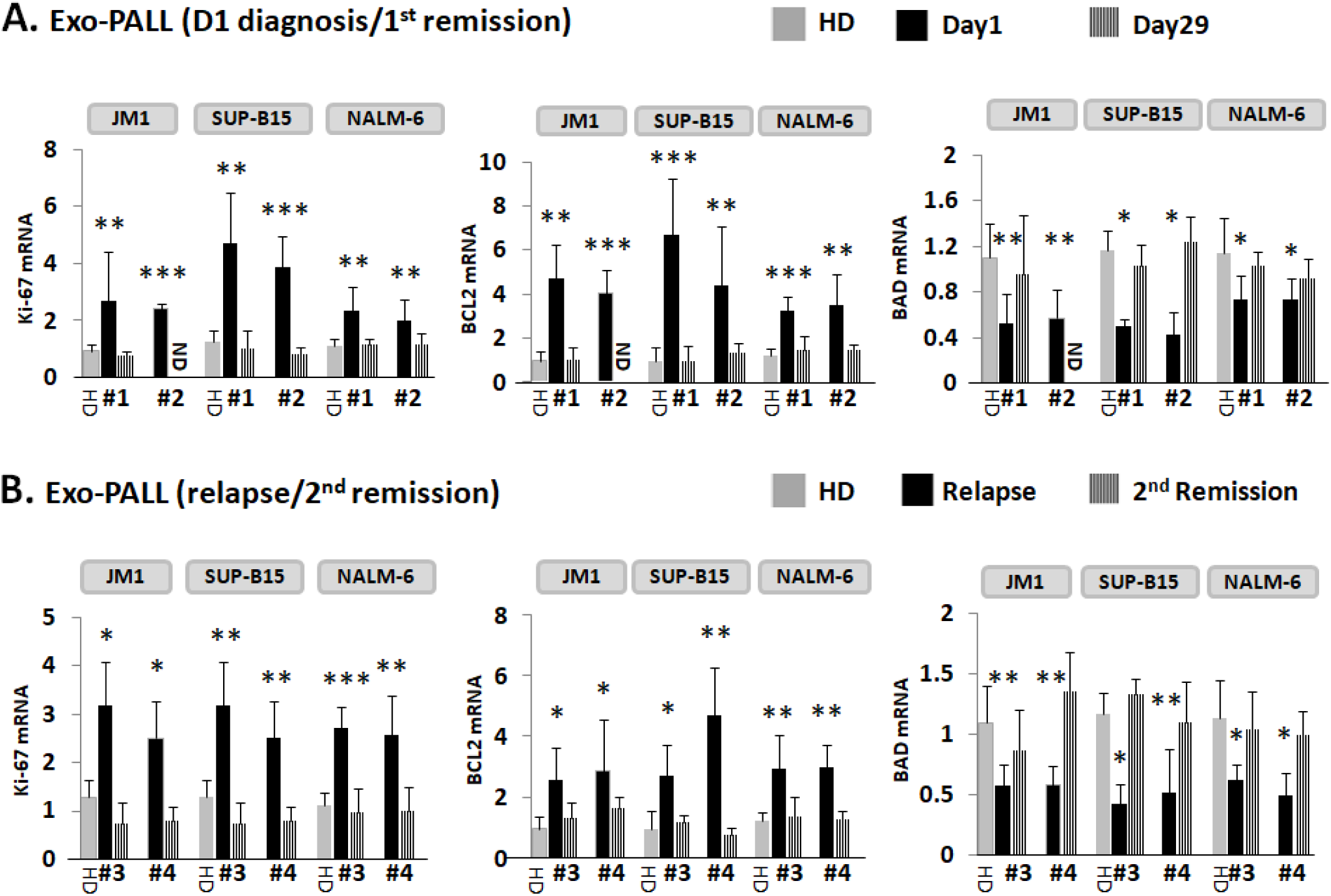
Exo-PALL treatment contributes to gene regulation in leukemia cell lines which correlates with disease stage. Three different cell lines were exposed with indicated Exo-PALL isolated from serum of pt#1 (Day1 diagnosis), pt#2 (Day29 1^st^ remission), pt#3 (first relapse) and pt#4 (2^nd^ remission). mRNA expression was evaluated by q-PCR. **(A)** Exo-PALL pt #1 up-regulates Ki-67 and BCL2 mRNA expression and down-regulate BAD mRNA expression in JM1, SUP-B15 and NALM-6 leukemia cell lines compared to HD exosomes. This augmentation effect of Exo-PALL on mRNA expression is no longer detectable during remission (pt#2; Day 29-remission). **(B)** Similar effect is obtained with Exo-PALL pt #3 and #4 at first relapse and in 2^nd^ remission. *(Pt#1, #2, #3, and #4: Exo-serum samples of four different patients. HD: healthy donor control serum. P value * p<0.05, **p<0.01, ***p<0.001)*

### Exo-miR expression by utilizing human cancer pathway finder array and validation of exosomal miR-181a by q-PCR

Exosomes from both CM of cell lines and serum samples (PALL and HD) were analyzed for their miRNAs array **(Fig. 5)**. We chose the Human Cancer Pathway Finder Array which allowed us to screen for 84 different miRNA commonly found in cancer. We found that exo-miR-181a-5p expression was 154-fold higher in JM1 cells **(Fig. 5A)**. Exo-miR-181a-5p was 40-fold higher in exosomes of PALL D1 (at day 1 of diagnosis, Pt#1) **(Fig. 5B)** but its expression reverted back to almost normal once the patient achieved remission at Day 29 (PALL D29 Pt#1) **(Fig. 5C)**. When comparing exo-PALL D29 vs. exo-PALL D1 in samples derived from the same patient (Pt#1), miR-181a-5p expression was 11-fold down regulated in the Day 29 samples compared to D1 **(Fig. 5D)**. We validated exo-miR-181a-5p expression in ALL cell lines and multiple PALL patients by q-PCR. miR-181a in Exo-JM1 was 140-fold and in Exo-SUP-B15 was 30-fold higher relative to control HD serum exosomes **(Fig. 5E)**. Exo-miR-181a-5p was also screened in 5 different PALL patients and data showed 3 to 5-fold higher expression of miR-181a-5p **(Fig. 5F)**. In serum exosomes of PALL patient #1, miR-181a-5p level at D1 of diagnosis was 20-fold higher compared to healthy donor, while at D29 (1 ^st^ remission) was only 3-fold higher compared to the control **(Fig. 5G 1**^**st**^ **graph)**. Similarly, in PALL Pt#2, miR-181a-5p expression in serum exosomes at D1 of diagnosis was higher (5-fold) than in HD, while at D29 (remission) went down to 0.15-fold compared to the control. **(Fig. 5G 2**^**nd**^ **graph)**. When comparing relapse vs. 2^nd^ remission, serum exosome miR-181a-5p expression in PALL Pt # 5 was 42-fold higher during relapse compared HD and decreased during 2^nd^ remission, although still remained 6-fold higher than the healthy control **(Fig. 5G 3**^**rd**^ **graph)**. Taken together, our data indicate that PALL D1 or relapse–derived exosomes are loaded with miR-181a-5p at the time of active disease status. Based on these, we hypothesized that leukemic exosomal miR-181a-5p could play a role in leukemia cell proliferation.

**Figure 5.**
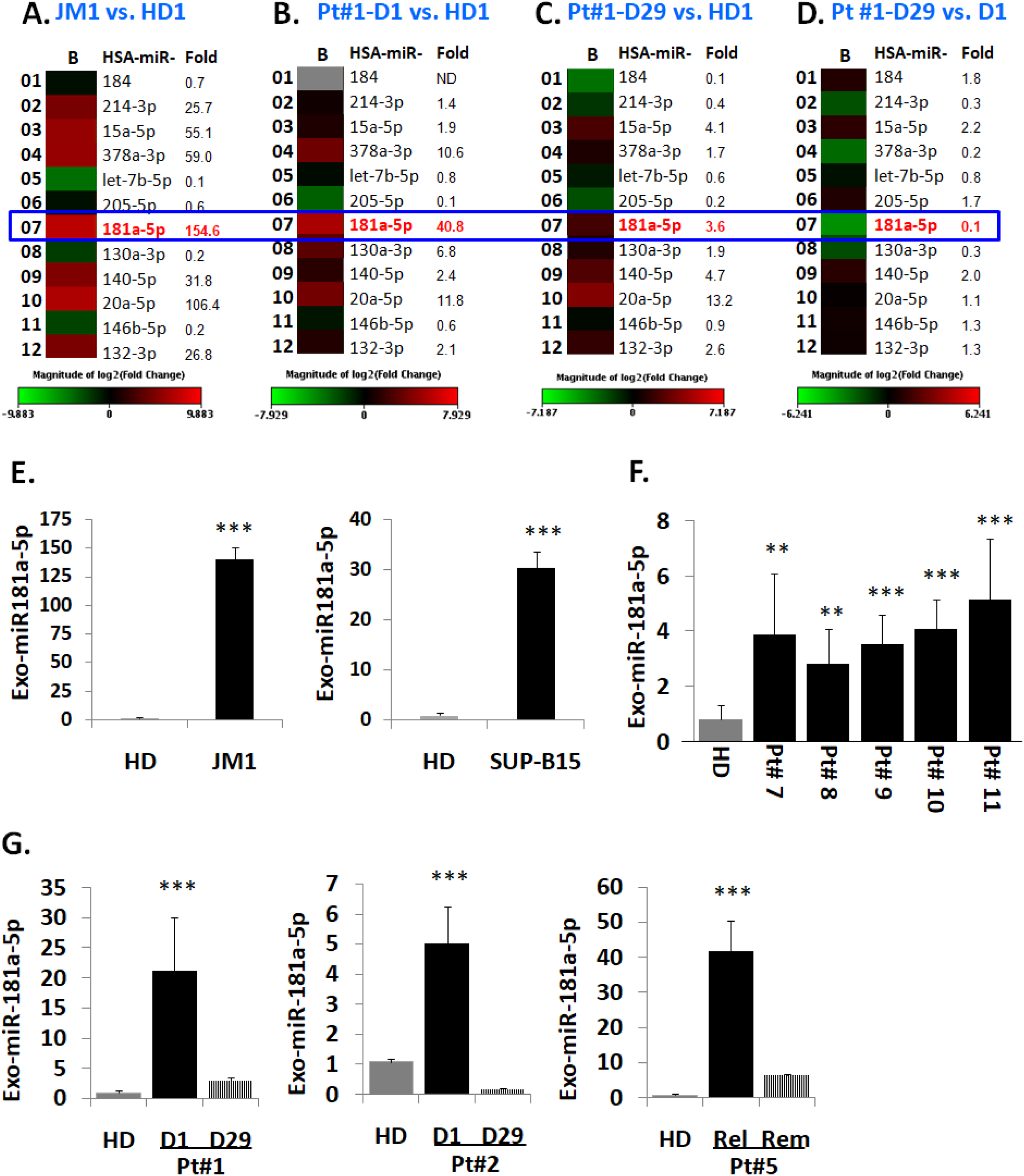
Heat map analysis of Exo-miR array by human cancer pathway finder and validation of exosomal miR-181a expression by q-PCR. Representative example of a heat map analysis shows differential miRNA expression between two groups overlaid onto PCR Human Cancer Pathway Finder Array plates show. (**A**) miR-181a is 154fold up-regulated in exosomes of JM1 cell line compared to HD exosomes; **(B)** miR-181a is 40 fold up-regulated in Exo-PALL of Pt# 1 at diagnosis (D1) compared to HD. **(C)** miR-181a is only 3.6 fold high in the same Exo-PALL (Pt#1) in remission (D29) compared to HD. **(D)** miR-181a is at baseline in Exo-PALL (Pt#1) upon remission (D29) compared to diagnosis (D1). **(E)** Expression of miR-181a-5p by q-PCR in Exo-CM of ALL cell lines JM1, and SUP-B15 compared to HD. **(F)** Exo-miR-181a expression in five PALL patients’ (Pt# 7, 8, 9, 10 and 11) samples by q-PCR compared to HD. **(G)** Expression of miR-181a-5p in Exo-PALL patients at diagnosis (D1), remission (D29), first relapse (relapse) and 2nd remission (remission) in three different ALL patient samples (Pt# 1, Pt# 2, & Pt# 5).*(Data represented are mean of triplicates. P value *p<0.05, **p<0.01, ***p<0.001).*

### Transfection efficiency of TexRed-siRNA into the target cell and dose optimization of miR-181a-5p inhibitor

To support the idea that miR-181a is a major player in leukemia cell proliferation, we explored miR-181a inhibition by a miScript miRNA inhibitor to reverse the induced cell proliferation. We first established and determined uptake-efficiency of a control siRNA by exosomes, by TexRed as per manufacturer recommendations **(Fig. 6)**. We transfected and loaded Exo-JM1 with TexRed-siRNA and co-cultured with JM1 cells. After for 24 hours, JM1 cells were harvested and analyzed for TexRed-siRNA uptake by flow cytometer. We observed around 50-60% JM1 cells were TexRed positive **(Fig. 6A)**. Once siRNA uptake was confirmed, Exo-JM1 was transfected and loaded with a specific miR-181a inhibitor and the exosomal miR-181a level was determined by q-PCR. Results showed that transfection of exosomes with an inhibitor resulted in a 70-88% silencing of exosomal miR-181a **(Fig. 6B)**. To rule out off-target activity causing non-specific inhibition, we randomly amplified miR-378 as an off-target for the miR-181a siRNA inhibitor. We found that there is no effect of the miR-181a inhibitor on miR-378 expression in exosomes, confirming specificity of miR-181a inhibitor **(Fig. 6C)**.

**Figure 6.**
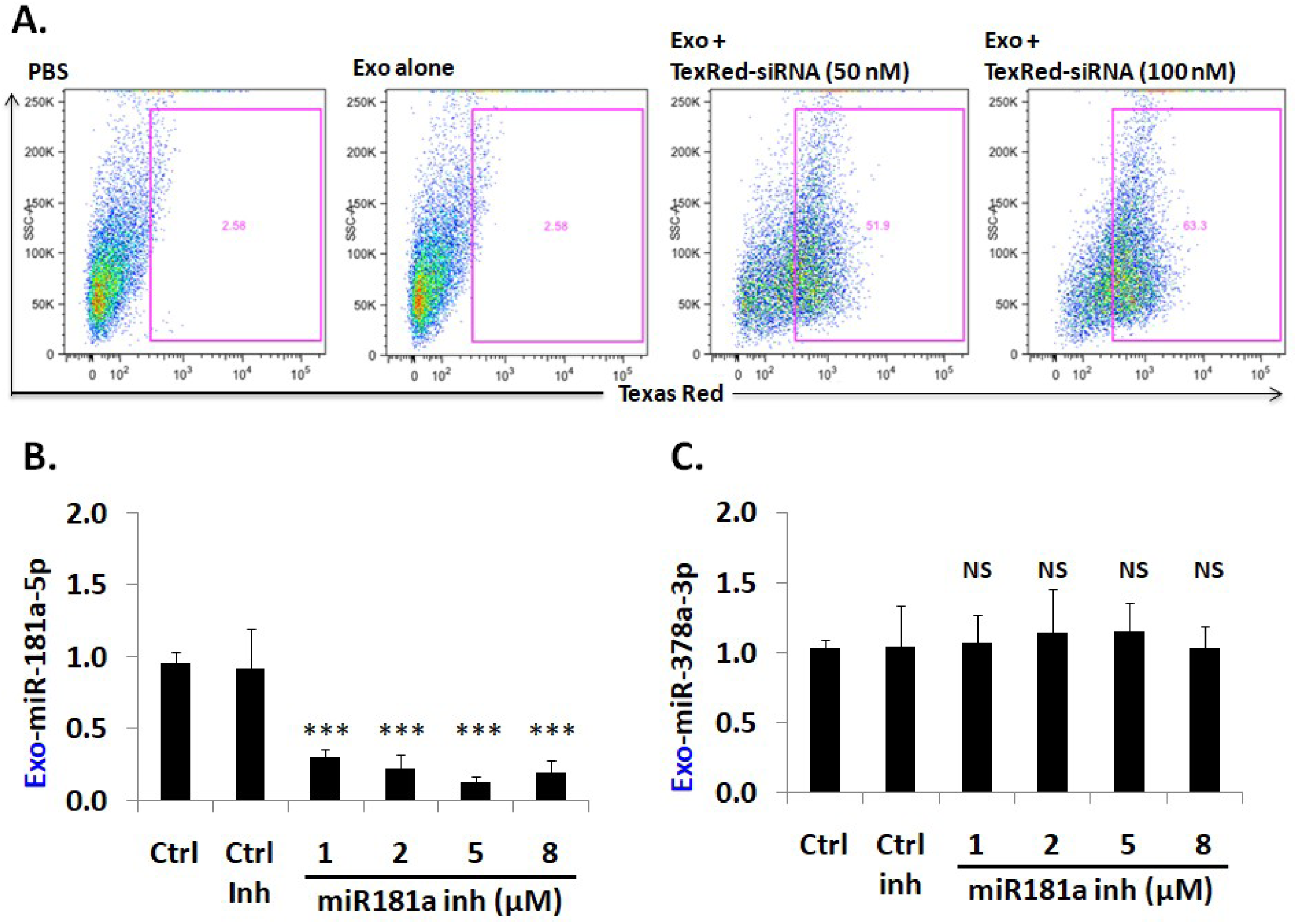
Cellular uptake efficiency of TexRed-siRNA and dose optimization of miR-181a-5p inhibitor. **(A)** TaxRed-siRNA transfection efficiency into Exo-JM1 and cellular uptake of exosome was measured by flow cytometer. FACS data show that 50-60% JM1 cells are TexRed positive compared with PBS/Exo alone control group. **(B)** Exosome transfection with Exo-miR-181a-5p inhibitor for miR-181a-5p silencing. Exo-JM1 were transfected with 1, 2, 5, 8 µM of miR-181a inhibitor. Exo-miR-181a-5p expression was carried out by q-PCR. Transfection of exosomes with an inhibitor resulted 70-88% silencing of exosomal miR-181a. **(C)** An unrelated miR-378 was amplified from the same cDNA by q-PCR. There is no effect of miR-181a inhibitor on miR-378 expression confirming specificity. *(SNORD61 was used as endogenous miR. P value *p<0.05, **p<0.01, ***p<0.001. NS: not significant).*

### PALL-derived exosomal silencing of miR-181a by miR-181a inhibitor reverses exosome induced cell proliferation

We hypothesized that PALL exosome-induced proliferation could be reversed by silencing of Exo-miR-181a by a specific miR-181a inhibitor **(Fig. 7)**. To explore further the role of miR-181a in cell proliferation, we treated leukemia cell lines for 24 hours with Exo-JM1 that were prior transfected with a miR-181a inhibitor in following conditions: PBS control (a), Exo-JM1 alone (b), Exo-JM1 + ctrl inhibitor (c), and Exo-JM1 + miR-181a inhibitor (d). Induced cell proliferation was observed after treatment with Exo-JM1 alone **(b)** or with Exo-JM1 + control inhibitor **(c)**, while in the Exo-JM1 + miR-181a inhibitor treated group **(d)**, cell proliferation remained at baseline, similar to PBS-treated cells **(a)**. We repeated these observations in five different cell lines (JM1, SUP-B15, REH, NALM6, and CL-01). All five cell lines showed a reproducible similar pattern of exosome-induced cell proliferation and this effect was reversed in the presence of the exosomal miR-181a-5p inhibitor **(Fig. 7A)**. These results suggest that miR-181a is functionally active in leukemia exosomes and contributing to induction of cell proliferation.

**Figure 7.**
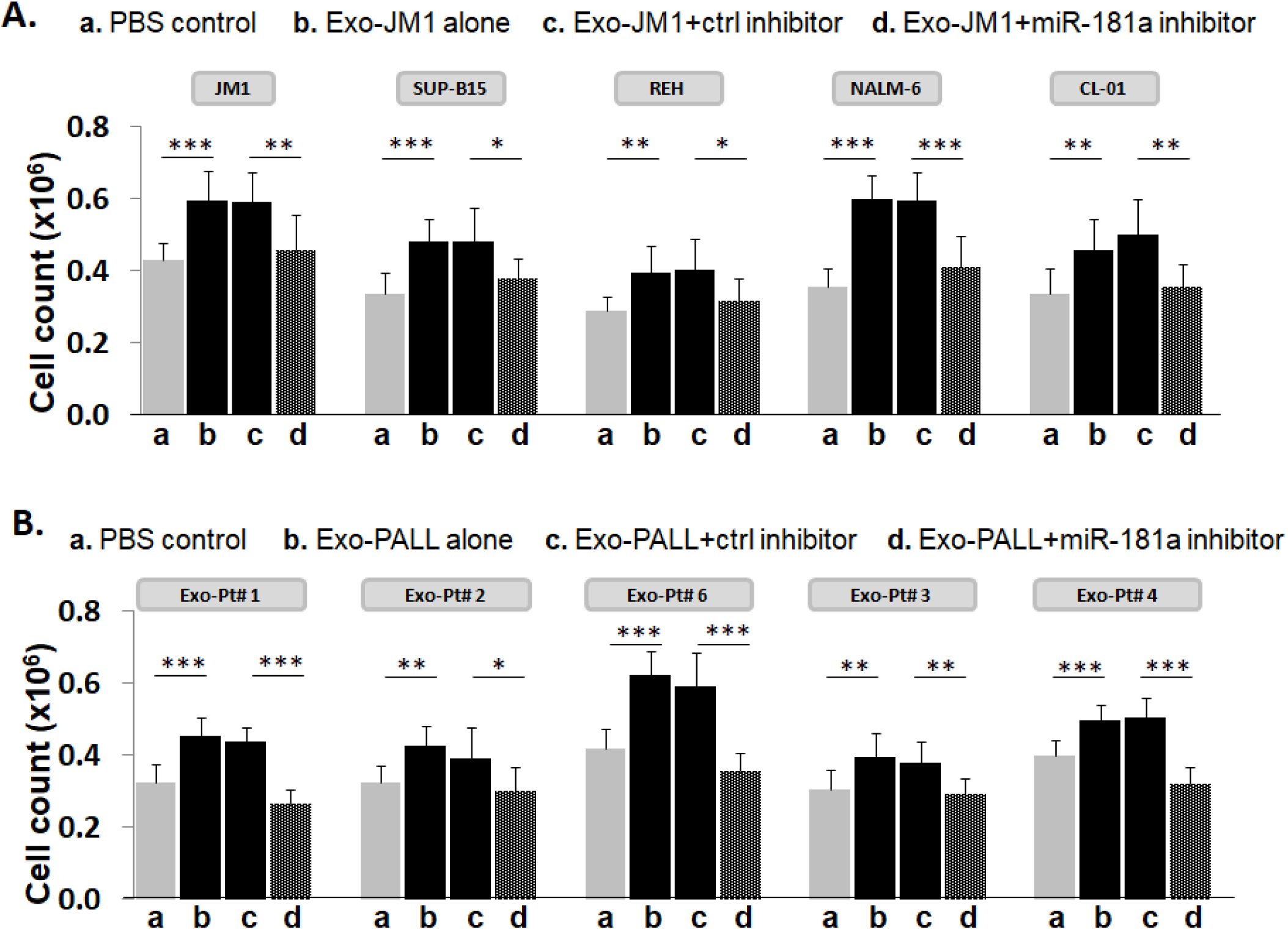
Leukemia derived-exosomal miR-181a silencing reverses exosome-induced cell proliferation. **(A) *Silencing of miR-181a in Exo-JM1 (CM) impairs cell proliferation:*** Indicated cells were treated with a) PBS, b) Exo-JM1, c) Exo-JM1 + control inhibitor, d) Exo-JM1 + miR-181a inhibitor for cell proliferation assay. Cell proliferation was observed after treatment with wild type Exo-JM1 **(b)** or with Exo-JM1+ control inhibitor **(c)** while in the Exo-JM1 + miR-181a inhibitor treated group **(d)**, cell proliferation remained at baseline, similar to PBS-treated cells **(a)**. These experiments were repeated in five different cell lines: JM1, SUP-B15, REH, NALM-6, and CL-01 as indicated**. *(B) Silencing of miR-181a in Exo-PALL (serum) impairs cell proliferation:*** JM1 cells were treated with a) PBS, b) Exo-PALL, c) Exo-PALL + control inhibitor, d) Exo-PALL + miR-181a inhibitor in cell proliferation assay. Cell proliferation was observed after treatment with Exo-PALL **(b)** or with Exo-PALL+ control inhibitor **(c)** while in the Exo-PALL + miR-181a inhibitor treated group **(d)**, cell proliferation remained at baseline, similar to PBS-treated cells **(a)**. Exo-PALL was derived from five different P-ALL patient samples (Pt#1, 2, 3, 4 and 6). *(P-value *p<0.05, **p<0.01, ***p<0.001).*

To confirm proof of concept, we treated JM1 cells with exosomes derived from five different PALL patient samples with or without prior miR-181a silencing by an inhibitor in the cell proliferation assay **(Fig. 7B)**. We observed, Exo-PALL alone **(b)** and Exo-PALL+ ctrl inhibitor **(c)** was able to induce cell proliferation compared to PBS control **(a)** while Exo-PALL + miR-181a inhibitor treated group **(d)** could not induce cell proliferation **(Fig. 7B)**. We obtained consistent outcomes in all five Exo-PALL samples. These results suggest that Exo-miR-181a from PALL patients’ serum contribute to leukemia cell proliferation, similarly to the results obtained from CM exosomes. Overall, our data suggest that exosomal miR-181a is a biologically active player with a functional role in leukemia cells that leads to induced cell proliferation.

## Discussion

In this study, we identified that exosomes induced cell proliferation in ALL B-lymphocytes as well as control B-lymphocytes. Furthermore, we showed that the cell proliferation is promoted by exosomal-miR-181a and reversible by mi-R181a silencing.

Characterization of exosomes was evaluated by Western blot and NanoSight analysis^31-32^. We titrated and optimized the dose of exosomes (100-500 µg/ml) for induction of cell proliferation. We show that 100 ug/ml and 250 ug/ml exosome treatment induces cell proliferation dose dependently, but at higher doses we observed a plateau effect. Literature illuminates that exosome dose optimization, for functional studies, varies from cell type to cell type. For example, in gastric cancer an exosomal dose of 50-400 µg/ml shows a dose-dependent cell proliferation^33^. Exosomes derived from mesenchymal cells showed a different dose & time-dependent angiogenesis in breast cancer^34^.

Here we report that exosomes derived from conditioned media (CM) of ALL cell lines and pediatric ALL patient’s serum induced cell proliferation in both leukemia and control B cells, augmented proliferative/pro-survival genes and downregulated pro-apoptotic genes. Chronic myeloid leukemia (CML) cell line LAMA84-derived exosomes promote proliferation genes (BCL-xl, BCL-w, survivin) and down regulates pro-apoptotic genes such as BAD, BAX, and PUMA^35^. However, there are no reports in literature on *acute lymphocytic leukemia*-derived exosomes induced cell proliferation.

To explore the physiological mechanism behind our results, we screened 84 different exo-miRs by use of the Human Cancer Pathway finder PCR array. We found that miR-181a was uniquely and significantly augmented in exosomes derived from P-ALL serum and leukemia cell lines. Furthermore, we showed that inhibition of exosomal miR-181a curbed the exosome induced cell proliferation. Based on our study, we suggest that exo-miR-181a might be responsible for induction of cell proliferation. While reports exist on miR-181a and cell proliferation, this is the first report delineating the effect of ALL-derived exosomes and exosomal miR-181a on cell proliferation. Signaling of miR-181a involves target genes of Wnt-signaling which lead to cell proliferation in ALL, by inhibiting WIFI signaling^36^. Verduci et al. reported that elevated miR-181a enhances cell proliferation in acute lymphoblastic leukemia by targeting EGR1^37^. In solid tumors, miR-181a has been described to play an important role for cancer progression specifically induction of cell proliferation, i.e. augmented miR-181a expression promotes prostate cancer cell proliferation by down-regulating DAX-1 expression and up-regulating PSA, CDK1, CDK2^38^. Over-expression of miR-181a leads to increased viability of osteosarcoma cells, inducing cell proliferation by augmenting BCL2, MMP9 and apoptosis was inhibited by down-regulation of p21 and TIMP3 expression^39^. In addition, augmented expression of miR-181a promotes cell proliferation and inhibits apoptosis in gastric cancer by targeting MTMR3^40^, and induced over-expression of miR-181a promotes cells proliferation in gastric cancer via the RASSF6-MAPK pathway^41^. Our novel data on the effect of PALL-derived exosomal miR-181a silencing on cell proliferation suggest that augmented miR-181a plays a role in lymphocytic leukemia cell proliferation via exosomal cellular uptake.

Silencing of miR-181a specifically at exosomal level was challenging. We optimized exosome transfection efficiency and cellular uptake of exosomal contents into the target cells. First, we established that exosome uptake efficiency into the targets cells is ∼50 - 65% for Texas-Red conjugated-siRNA, similar to what others have reported for TexRed labeled exosomes in HuH-7 cells^42^, and alexa-flour-488-conjugated siRNA labeled exosomes into the HTB-177 cells^43^. Exosome uptake and cargo delivery into target cells involves multiple mechanisms, such as soluble signaling, juxtacrine signaling, membrane fusion, phagocytosis, macro-pinocytosis, and receptor/raft mediated endocytosis^44^. We also showed that successful transfection with a miR-181a inhibitor into exosomes resulted in ∼ 80 - 90% inhibition of exo-miR-181a expression. The miR-181a inhibitor used in this experiment was highly specific to miR-181a since it did not silence the expression of another randomly selected miR, namely miR-378a. Other groups have attempted exosomal silencing by siRNA with similar results; i.e. exosomal delivery of a PLK-1 gene-specific siRNA silencer in UMUC3 bladder cancer cell lines showed >60% silencing of PLK-1 expression^45^. Endo et al. showed lipid nanoparticle as delivery system carrying siRNA (90% encapsulation of siRNA into the lipid nanoparticle) for the silencing of indoleamine2, 3 dioxygenase 1 gene which has implication for cancer immunotherapy^46^.

Exosome-based siRNA-inhibitor delivery can be considered a better vehicle system than liposomal-based or other vector delivery systems for several reasons: 1. Exosomes are biological nano-carrier systems that easily communicate with target cells for exosomal cargo delivery. 2. Exosomes harvested from the patient’s own body fluids or cell culture conditioned medium are non-immunogenic and can be utilized for autologous loading of a biological agent. 3. Exosomes are considered non-toxic in contrast to other transfection reagents^4^. Our data supports that miR-181a silencing *in exosomes* reverses exosome-induced cell proliferation. Although some reports in literature educate that *cellular inhibition* of miR-181a suppresses cell growth and proliferation^36, 41^, none have reported the effect of *exosomal* miR-181a silencing upon cell proliferation in ALL.

## Conclusions

Exo-miR-181a expression is abundantly up-regulated in exosomes derived from P-ALL patients’ serum and CM of ALL cell lines and correlates with leukemia cell proliferation; subsequent silencing of Exo-miR-181a reverses exosome-induced leukemia cell proliferation. Therefore, exosomal miR-181a inhibition can act as a novel synergistic target for growth-suppression in pediatric acute lymphocytic leukemia.

## Methods

### Cell lines and cell culture

ALL cell lines were purchased from ATCC (SUP-B15, JM1 and Nalm-6/CRL-1929 ™, CRL-10423™and CRL-3273™) and REH cells from DSMZ (Cat# ACC 22). CL-01 was donated by the Chiorazzi lab, FIMR, NY. Exosomes were isolated by ultracentrifugation from conditioned medium of cell lines (Exo-CM)^30^. miRNA silencing inhibitors: miScript inhibitor negative control (20 nmol, Qiagen, cat# 1027272), anti-hsa-miR-181a-5p miScript miRNA Inhibitor (20 nmol, Qiagen, cat# MIN0000256).

### Serum samples

PALL patients (n=11) and healthy donor (HD) (n=4) serum samples were obtained after informed consent according to an IRB approved protocol in accordance with the Helsinki Declaration ***(Supplemental Table 1)***. P-ALL serum samples were collected at different leukemia disease stages: new diagnosis (Day 1), remission (Day29), relapse or second remission. Exosomes were isolated from serum by ultracentrifugation as described (Exo-PALL)^30^.

### Depletion of exosomes from the FBS

To avoid contamination by exogenous exosomes contained in the fetal bovine serum (FBS) that is added to culture medium, FBS was converted into exosome-free FBS by ultracentrifugation.^30^ Briefly, FBS was subjected to centrifugation for 10 min at 300 x*g*. Supernatant was collected and centrifuged for another 10 min at 2000 x*g*. Again, supernatant was collected and centrifuged for 30 min at 10,000 x*g*. Then supernatant was ultra-centrifuged overnight at 100,000 x*g*. The following day, FBS supernatant was collected and labeled as exo-free FBS. Each centrifugation step was carried out at 4°C.

### Cell culture set up for exosome production

Each cell line was cultured in exo-free FBS/exosome depleted FBS cell culture medium when cells were expanded for the harvest of conditioned medium (CM) exosomes. Cells were plated (1×10^6^/ml) in 10 ml total volume in a 100mm tissue culture dish. After 48 hours, CM of each cell line was harvested by centrifugation and filtered through a 0.22 micron filter (Millipore) to remove cell debris. Filtered CM was used for exosome isolation by ultracentrifugation.

### Exosome isolation

Exosomes were purified by ultracentrifugation method^30^. In brief, human serum and CM from cell lines were subjected to centrifugation for 10min at 300 x*g*. Supernatant was collected and centrifuged for 10min at 2000 x*g.* Again, supernatant was collected and centrifuged for 30min at 10,000 x*g*. Then supernatant was ultra-centrifuged for 120min at 100,000 x*g.* Pellets were re-suspended in PBS and again ultra-centrifuged for 120min at 100,000 x*g.* Pellets containing exosomes were harvested and reconstituted in 250 μl PBS. Each centrifugation step was carried out at 4°C. The protein content of the exosomes was measured using a BCA protein assay kit (Bio-Rad). Exosomes were aliquoted and stored at −80°C for further usage.

### Exosomal CD63 and CD81 expression by flow cytometer

We used exosome isolation and analysis kit (cat: ab239682) from Abcam and followed protocol as directed. Briefly, Exosomes (10 µg) was taken into 50 µl PBS. Captured beads (100 µl) were mixed with 50.0 µl of exosomes in cytometer tube. Mixture was incubated at RT in dark overnight. Primary detection antibody (CD81-biotin conjugated): 5.0 µl of primary detection antibody (CD81) were added and incubated at 2-8 °C for 1.0 hour in dark. Washing steps: 1.0 ml of assay buffer was added to samples, mixed by tapping and centrifuged at 2000 rpm for 5.0 minutes. Supernatant were decanted and sample/pellet were reconstituted in 100 µl assay buffer. Secondary detection reagent (streptavidin-PE conjugated): 5.0 µl of secondary antibody (PE-conjugated) were added in each tube and incubated at 2-8 °C for 30 minutes in dark. Then washing step was repeated. Samples/pellet was reconstituted in 350 µl assay buffer and read at Fortessa.

### Exosome induced-cell proliferation assay

JM1, SUP-B15 and NALM-6 leukemia cell lines (0.2×10^6^/well) were seeded in 96-well plates and cultured in exo-free FBS medium. Each well was loaded with exosomes (250 µg/ml) and incubated at 37 °C. After 24 hours, cells were mixed by gently pipetting. Cell proliferation was analyzed by live cell counting using trypan blue/hemocytometer method under the microscope.

### Cellular RNA extraction and cDNA preparation

Total RNA was extracted from SUP-B15, JM1, or CL-01 cells with the use of a Trizol reagent (Invitrogen) method. Quality and quantity of RNA was analyzed by NanoDrop ND1000 spectrophotometer. Around 2 to 5 µg of total RNA was used for cDNA synthesis by q-PCR. Oligo-dT primers (Invitrogen) and M-MLV reverse transcriptase (cat # 28025-013, Invitrogen) were used for cDNA synthesis as per manufacturer’s protocol.

### Cellular mRNA expression by q-PCR

Primers sequences, probe numbers and gene accession numbers from universal probe library (UPL) of Roche Applied Science are described ***(Supplemental Table 2)***. Gene amplification was carried out by q-PCR with cDNA as template and Eurogentec master mix. Fold change was calculated by comparing treated vs. untreated groups. Data were analyzed with RQ manager version 1.2.1 (Applied Biosystems, Foster City, CA). The data are expressed as fold change. GAPDH as an endogenous reference gene.Exo-JM1 and Exo-SUP-B15 were loaded on JM1 cells. After 24 hours of exosomes loading, cells were harvested and mRNA extracted for q-PCR analysis. We amplified proliferative genes (Ki-67, PCNA), pro-survival (MCL1 and BCL2) and pro-apoptotic pathways (BAD and BAX) after incubation with exosomes. We also tested exosomes purified from HD and four P-ALL patients at different disease stage: P-ALL D1 (day 1 of diagnosis), and P-ALL D29 (1^st^ remission) (pt #1 and pt #2), P-ALL relapse, P-ALL 2^nd^ remission (pt #3 and pt #4) in three leukemia cell lines.

### Exosomal RNA isolation

CM was harvested from JM1 and SUP-B15 (2 x 10^6^ cells/ml) after 48hr and exosomes were isolated by ultracentrifugation (10min at 300x*g*, 10min at 2000 x*g*, and30min at 10,000 x*g)*. Collected supernatant was processed for isolation of exosomal RNA (ExoRNeasy Kit (MAXI) - Qiagen). Similarly, exosomal RNA from human serum samples was extracted utilizing the ExoRNeasy Kit (MIDI) (cat # 77044).RNA was quantitated by NanoDrop.

### Exosomal miR-181a-5p expression by PCR

Exo-RNA was converted into cDNA by miScript II RT kit (Cat # 218161, Qiagen). Expression of miRNA was evaluated by q-PCR using a miScript SYBR@Green PCR kit (Cat # 218073, Qiagen).Screening of miRNA expression for 84 different miRNAs was performed on Exo-RNA isolated of CM cell lines and serum exosomes, using a Human Cancer Pathway Finder miScript miRNA PCR Array(Cat # MIHS-102ZF, 331221-Qiagen). Expression of miR-181a was validated by q-PCR using primers (cat # MS00008827, Qiagen).

### Exosomal miR-181a silencing with a specific miR-181a inhibitor

#### A. Confirmation of Exosomal uptake of TexRed-siRNA and cellular uptake of exosomes

Exosomes were labeled with TexRed-siRNA (SBI System Bioscience) as per manufacturer’s recommendations. TexRed-labeled JM1 exosomes were then co-cultured with JM1 cells for 24 hours. Presence of TexRed-siRNA (introduced in target cells by exosomes) into the JM1 cells was assessed by flow cytometer.

#### B. Dose titration of miR-181a inhibitor

Exo-JM1 were transfected with different concentrations of the miR181a inhibitor (1, 2, 5, 8 µM) to determine optimal dosing. Then, RNA was isolated from transfected exosomes for cDNA synthesis and miR-181a expression by q-PCR.

#### C. Exosomal miR-181a silencing

300 µg of exosomes (either CM- or serum-derived) was transfected as follows: 1/ transfection reagents only;2/ transfection reagent and control inhibitor;3/ transfection reagent with miR-181a inhibitor (silencing). Each transfection mixture was co-cultured with JM1 cells for 24hrs allowing for exosomal uptake by the target cells. Cell count was carried out by light microscopy.

### Statistical Analysis

To compare the mean values between two groups, the unpaired t-test was used. Statistical significance was defined as p<0.05. All results are represented as Mean ± SD. Each experiment was performed in triplicates.

## Data Availability

The data generated during the current study are available from the corresponding author on a reasonable request.

## Acknowledgements

This work was supported by the Pediatric Oncology Fund at Staten Island University Hospital, The DiMartino Family Foundation and Island Auto group. We also thank Malvern Instruments for the characterization of exosomes by NTA.

## Authors Contributions

S.H. performed the experiments, designed the experiments, analyzed the data and wrote the manuscript. S.V. designed the experiment, reviewed and edited the manuscript, and provided all the required reagents.

## Abbreviations

P-ALL: Pediatric Acute Lymphoblastic Leukemia
Exo-PALL: Exosomes derived from Pediatric Acute Lymphoblastic Leukemia
Exo-HD: Exosomes derived from Healthy Donor
Exo-CM: Exosomes derived from Conditioned Medium

