## Supplemental file for "Silencing of Exosomal miR-181a reverses Pediatric Acute Lymphocytic Leukemia Cell Proliferation"

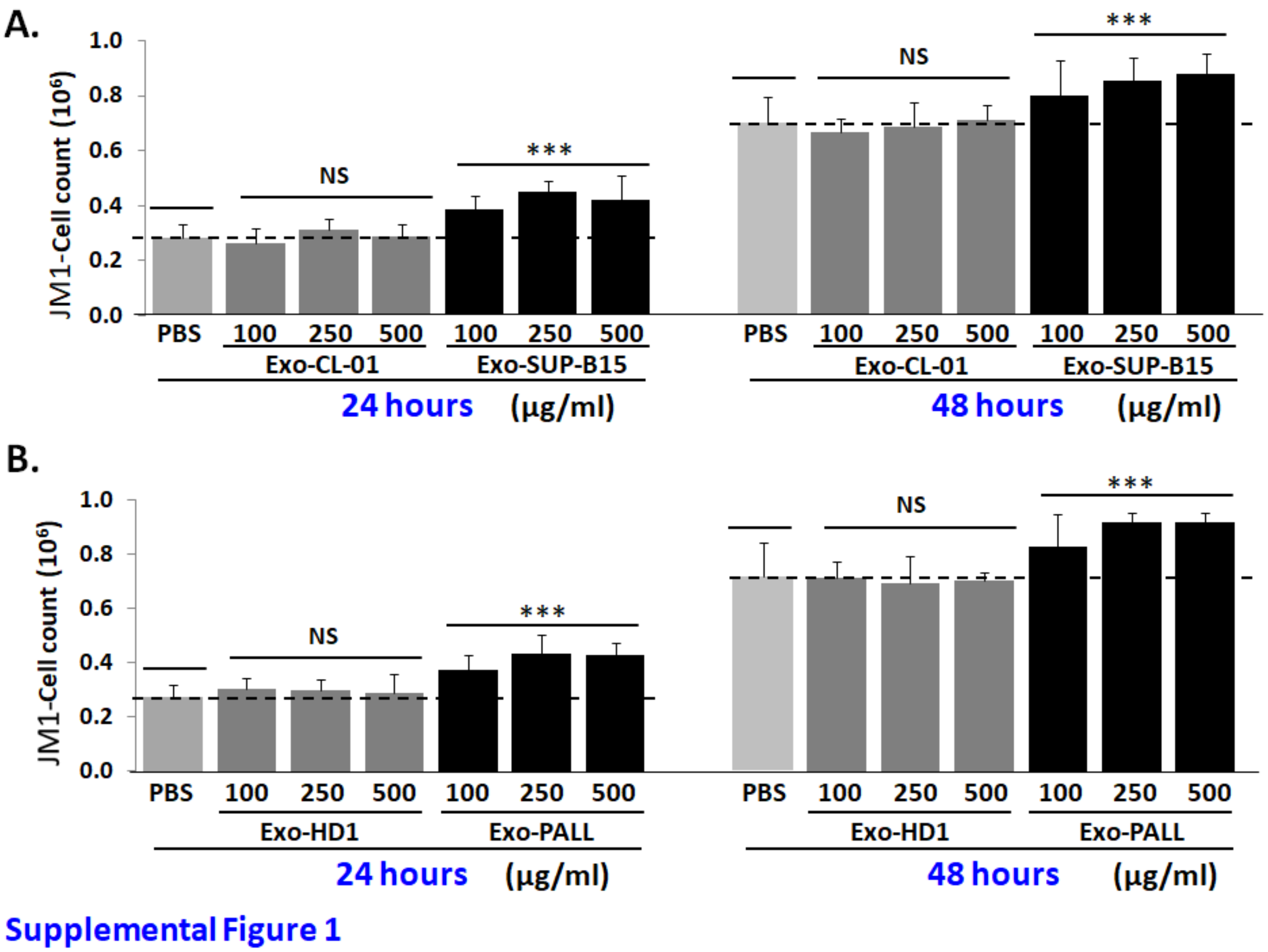


**Supplementary Figure 1**

**Dose titration and time kinetics of exosomes enhancing cell proliferation. (A)** JM1 cells were treated with Exo-CL-01 (normal B cell line) and Exo-SUP-B15 (leukemia B cell line) in three different dosages (100, 250, 500 µg/ml). Cells were counted at 24 hours and 48 hours after treatment. **(B)** JM1 cells were loaded with Exo-HD and Exo-PALL in three different dosages (100, 250, 500 µg/ml) in JM1 cells. Cells were counted at 24 hours and 48 hours after treatment. Exosomes induced cell proliferation was present at optimal concentration of 250 µg/ml of exosomes. *(P value * $p<0.05$, ** $p<0.01$, *** $p<0.001$. NS; not significant)*

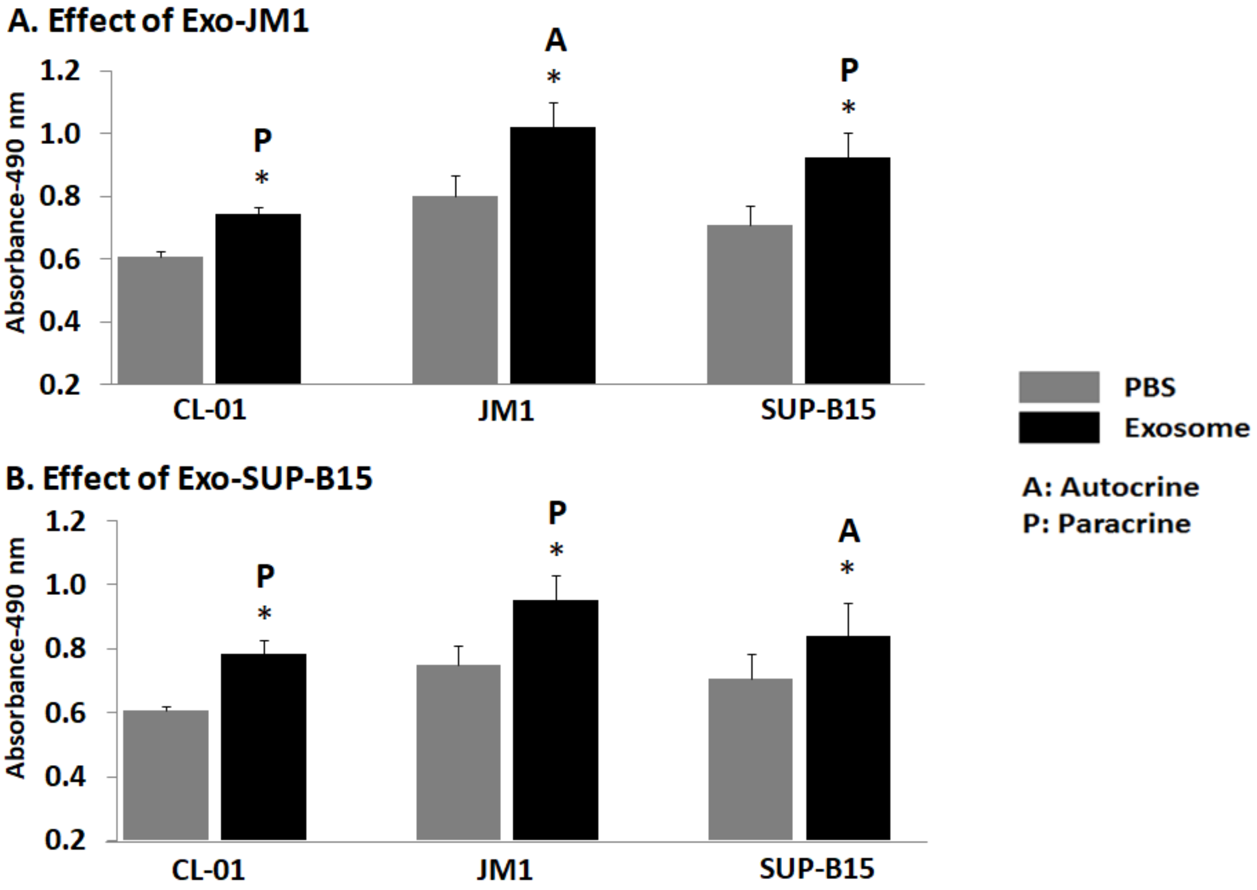


Supplemental Figure 2

**Supplementary Figure 2**

**Exosomes induced cell proliferation by MTS assay.** (A) CL-01, JM1, and SUP-B15 cells were plated (0.1x$10^6$/well) in quadruplets. Exo-JM1 (JM1 cell-derived exosomes) was loaded (250 µg/ml) on the CL-01, JM1, and SUP-B15 for 24 hours. Next day, MTS were added into the culture plate and plate was read at 490 nm. (B) CL-01, JM1, and SUP-B15 cells were plated (0.1x$10^6$/well) in quadruplets. Exo-SUP-B15 (SUP-B15 cell-derived exosomes) was loaded (250 µg/ml) on the CL-01, JM1, and SUP-B15 for 24 hours. Next day, cell proliferation was quantitated by MTS. Data was analyzed by PBS as control (P value * p<0.05, **p<0.01, ***p<0.001. NS; not significant).

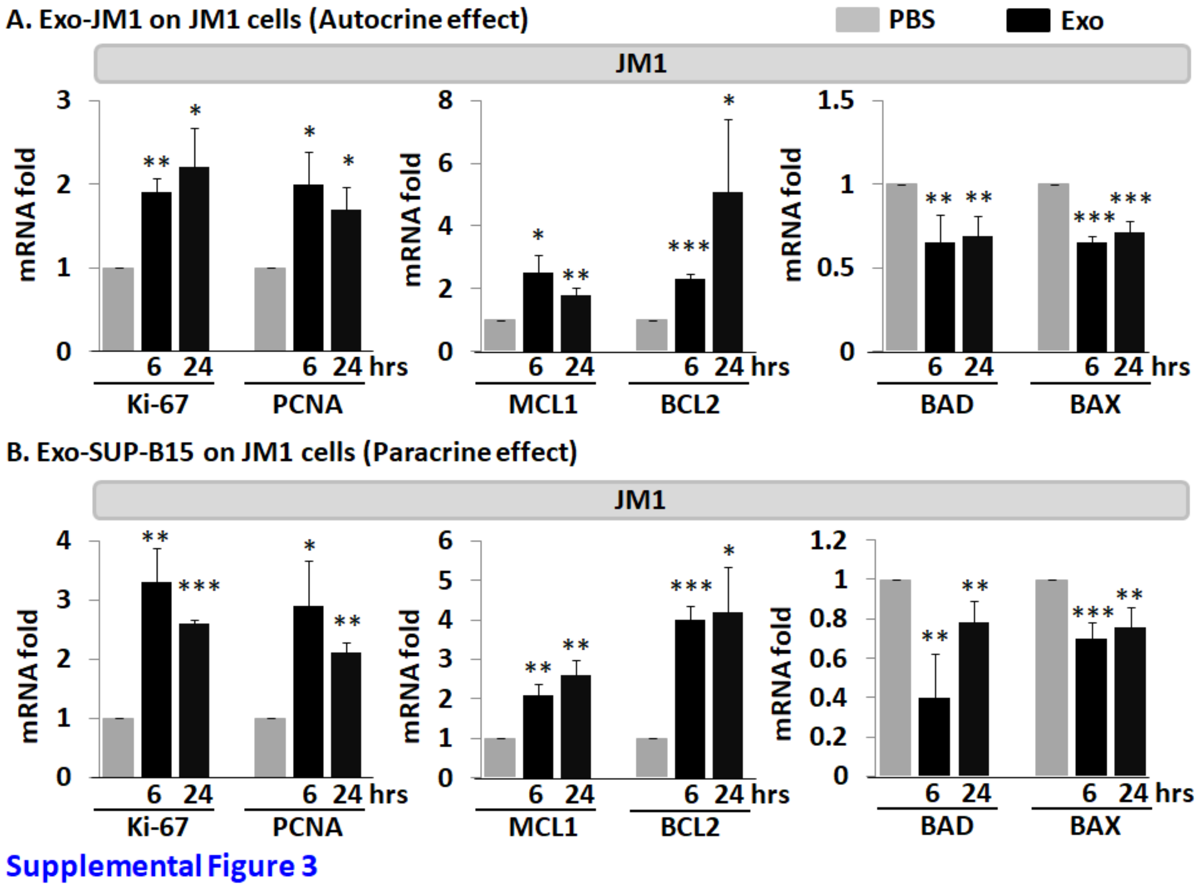


**Supplementary Figure 3**

**Exo-CM (JM1 and SUP-B15) regulates proliferative, pro-survival, and pro-apoptotic genes. (A)** JM1 cells were exposed with Exo-JM1 (250 µg/ml) and cultured cells were harvested for RNA isolation at 6 hours and 24 hours post treatment. Indicated mRNA (Ki-67, PCNA, MCL1, BCL2, BAD, BAX) expression analyzed by q-PCR. **(B)** JM1 cells were exposed with Exo-SUP-B15 (250 µg/ml). Cells were harvested for RNA isolation at 6 and 24 hours after treatment. Indicated mRNA (Ki-67, PCNA, MCL1, BCL2, BAD, BAX) expression analyzed by q-PCR. Data represented are mean of three experiments. *(Ctrl: PBS only/no exosomes- P value *p<0.05, **p<0.01, ***p<0.001).*

**Supplemental Table 1**

**List of healthy donors and PALL serum samples**

| Healthy Donor # | Serum Sample Code |
|---|---|
| 1 | HD77 |
| 2 | HD78 |
| 3 | HD79 |
| 4 | HD80 |
| PALL patient # | Serum Sample Code |
| 1 | PALL01 D1 |
| | PALL01 D29 |
| 2 | PALL02 D1 |
| | PALL02 D29 |
| 3 | PALL14c relapse |
| | PALL14c 2nd remission |
| 4 | PALL24 relapse |
| | PALL24 2nd remission |
| 5 | PALL14 relapse |
| | PALL14 2nd remission |
| 6 | PALL25 D1 |
| 7 | PALL03 |
| 8 | PALL04 |
| 9 | PALL05 |
| 10 | PALL05b |
| 11 | PALL14a |

**Supplemental Table 2**

**Human primers from universal probe library (UPL)**

| Genes | Accession # | Probe # | Primers sequences |
|---|---|---|---|
| PCNA | J04718.1 | 77 | For: 5'-CTTTTTCGCGCCAAAGTC-3’ |
| | | | Rev: 5‘-CTGCGGAAAAACCCTTGAT-3' |
| Ki-67 | NM_002417.4 | 53 | For: 5‘-CGCGTAAGTCAAGACCAAAAT-3' |
| | | | Rev: 5‘-GGTCAAGCTCTTGTTCAGGTG-3' |
| BAD | AF031523.1 | 45 | For: 5‘-ACCAGCAGCAGCCATCAT-3' |
| | | | Rev: 5‘-GGTAGGAGCTGTGGCGACT-3' |
| BAX | U19599.1 | 55 | For: 5‘-CAAGACCAGGGTGGTTGG-3' |
| | | | Rev: 5‘-CACTCCCGCCACAAAGAT-3' |
| MCL1 | AF118124. | 4 | For: 5‘-AAGCCAATGGGCAGGTCT-3' |
| | | | Rev: 5‘-TGTCCAGTTTCCGAAGCAT-3' |
| BCL2 | AY220759.1 | 23 | For: 5‘-TTGGTATCCTTCTCTTTCAGCAC-3’ |
| | | | Rev: 5‘-ATGGCATTGACGAAGAGGAT-3' |
| GAPDH | NM_002046.3 | 60 | For: 5’-AGCCACATCGCTCAGACAC-3’ |
| | | | Rev: 5’-GCCCAATACGACCAAATCC-3’ |
